# A causal approach to understanding management factors, reproductive underperformance and reproductive infections in dairy cattle

**DOI:** 10.1101/569764

**Authors:** Adis Softic, Kassahun Asmare, Erik Georg Granquist, Jacques Godfroid, Nihad Fejzic, Eystein Skjerve

## Abstract

The objective of this study was to investigate causal associations between cattle farms’ management practices and reproductive disorders (abortion, stillbirth, retention of placenta, metritis). Besides, direct causal associations between farms’ management and reproductive infections (*Chlamydia abortus, Coxiella burnetii*, and *Neospora caninum*), reproductive disorders and infections were also investigated in this study. As a secondary objective, constraints that affect the production in cattle farms were examined. The study was carried out in the north-western (Una-Sana), western (Canton 10) and central part (Central Bosnia Canton) of Bosnia and Herzegovina. A total of 201 farms were selected for participation. A semi-structured questionnaire-based interview was conducted among farmers/managers from January 1^st^ to August 31^st^, 2015. The 40 questions were divided into three groups: socio-demographic, management, and information on reproductive performances in cattle. Supplementary questions were asked about the perceived primary constraints of the production.

A multivariable mixed-effects logistic regression was used to screen management factors for potential statistical influence. All investigated outcomes were associated with farms’ management. The final multivariable models were merged into a Structural Equation Model (SEM). The causal model was then specified graphically. The SEM model showed that herds that experienced abortions (OR=4.3) and stillbirth (OR=6.7) were associated with *N. caninum* seropositivity. Also, herds that experienced retention of placenta were strongly associated with the occurrence of metritis (OR=10.1). *C. abortus* and *C. burnetii* herd seropositivities were mainly associated with environmental factors and contact with potential intermediate hosts.

Our study demonstrated that management practices on dairy farms in Bosnia and Herzegovina contributed to the occurrence of severe reproductive outcomes and reproductive infections. *N. caninum* seems to be an infectious agent that substantially contributed to the reproductive underperformance. Further we demonstrate the need for using causal models in understanding complex relationships.

## Introduction

Agriculture is one of the most vital sectors of economy for the countries in Western Balkan. Most of them have substantial share in the total state gross value added (GVA) ranging from 7.6% in Bosnia and Herzegovina (BH) to 20.1% in Albania. [1]. The situation in the agricultural sector is gradually improving in most of Western Balkan countries, but there are still some critical steps in reforming the agricultural policy and its harmonising with the EU standards. Agriculture is especially important for the economic development of BH [2]. The government of BH has committed to raising the little share (7.6%) of agriculture in the gross domestic product (GDP) through the focused strategic approach to the sector [2]. To promote this development, a new legislation was drafted to enforce compliance with standards set by the European Union (EU) for agricultural commodities. Livestock production is one of the focus areas, where the agriculture economy is expected to grow. Production of milk and dairy products increased in the period 2010-2015, with the production of approximately 701 million litres of milk in 2015 [3]. Also, authorization to the export of dairy products to the EU (in August 2016) represents an opportunity for further intensification of this production. This opportunity makes milk one of the principal export commodities of the country. To cope with the new requirement in the dairy production, infectious and non-infectious causes of the low reproductive performance of cattle become a priority to improve production performances.

Performance indicators are essential tools for monitoring overall reproductive performance in cattle. Some indicators are specific and easily recognised, such as abortion, retained placenta, metritis, and stillbirth. Problems in the postpartum period are usually well recognised by the farmers. Although a reporting system for serious reproductive failures (“Official Gazette of BH” issue number 4/16) is updated annually, it has not been fully implemented and is not made available to the farmers as a decision tool. The subtler effects of reproduction underperformance such as prolonged calving intervals, repeat breeder cows and losses due to early embryonic death require more systematic recording. These reproductive problems are usually calculated from the recorded reproductive events including calving date, insemination dates, and pregnancy outcome [4]. Fertility measures calculated from reproductive events can be divided into two categories as fertility scores and interval traits [5].

Losses due to infectious diseases are typically reflected in impaired (re)productivity and lack of market access. In a previous paper, a relatively high occurrence of exposure to infectious agents in cattle; *Chlamydia abortus, Coxiella burnetii*, and *Neospora caninum*, in selected cantons of BH [6] was demonstrated. Each of these pathogens can lead to clinical disease and have effects on bovine reproduction [7-9]. Notably, control measures against brucellosis in small ruminants seem to have more or less eliminated *Brucella melitensis* spill-over to cattle in the selected study regions, and there is no clear indication that *Brucella abortus* is present in cattle in BH [6]. Other infectious agents, like Bovine Viral Diarrhoea Virus (BVDV), may also be directly associated with reproductive failures [10, 11] and indirectly because transiently or persistently infected herds would likely lead to some immunosuppression of cattle [12]. Unfortunately, there are still knowledge gaps regarding the importance of BVDV infection in BH.

While we have documented high levels of certain reproduction pathogens in three cantons in BH, the importance of these agents remains unclear. Reproductive performances are also influenced by a multitude of management factors [13]. Cattle farming in BH still faces some limiting factors, and it is mainly taking place on small farms with an average area below 2 hectares and a high level of land fragmentation. Furthermore, there is a limited number of animals per farm (1-10) with approximately 75% of the farms having one or two cows [14]. Poor housing conditions followed by poor management linked to the inadequate education of the farmers may also contribute substantially to low productivity and losses in animal production. Management may also directly influence the occurrence of infectious agents. In this case, management will have an indirect effect on reproduction through increasing exposure to infectious agents. To be able to suggest sound interventions for improving reproductive performance, there is a need to evaluate which factors are dominant and their causal relationships with reproduction performance. As an improved production will also typically imply larger herds, new factors may also become more critical as, e.g., the dynamics of infections, that depend on herd size.

Although improvement of the cattle sector in BH partially depends on the views and motivation of the farmers/ owners, the scientific data from farms often do not include the perspective of the farmer/ owner. Importantly, their perceived main constraints on production will complement the data collected through epidemiological studies.

The primary objective of the present study was to describe the importance of management practices and to identify the causal interaction between management practices and selected infectious diseases (*C. abortus, C. burnetii*, and *N. caninum*) on reproductive underperformance and specific reproduction disorders. The secondary objective was to identify perceived constraints that affect production in the cattle farms in the study area.

## Materials and methods

### Study area and study design

The study was carried out in the north-western (Una-Sana Canton), western (Canton 10) and central (Central Bosnia Canton) parts of BH, which covers a 23.9% of the total BH territory. Based on a detectable herd-level prevalence of 5% and 95% level of confidence, a minimum sample size of 60 farms (total n=180) was selected in each region to achieve the detection of the agent. In the period from January 1^st^ to August 31^st^, 2015, a total of 201 cattle farms were visited, blood samples were collected, and a questionnaire-based interview was conducted. Four farms were excluded due to insufficient farm records, leaving 197 farms in the final dataset for statistical analyses (Fig 1). A complete farm sampling frame was unavailable, and farm selection was made using the available list of farms at the municipal level, within each region. In each farm, all female animals older than 12 months were selected for sampling. The breed composition among chosen farms was different, including; Simmental, Montafon, Holstein-Friesian, Red Angus, and their crosses. This study provided a total of 1970 individual blood samples.

**Fig 1.**
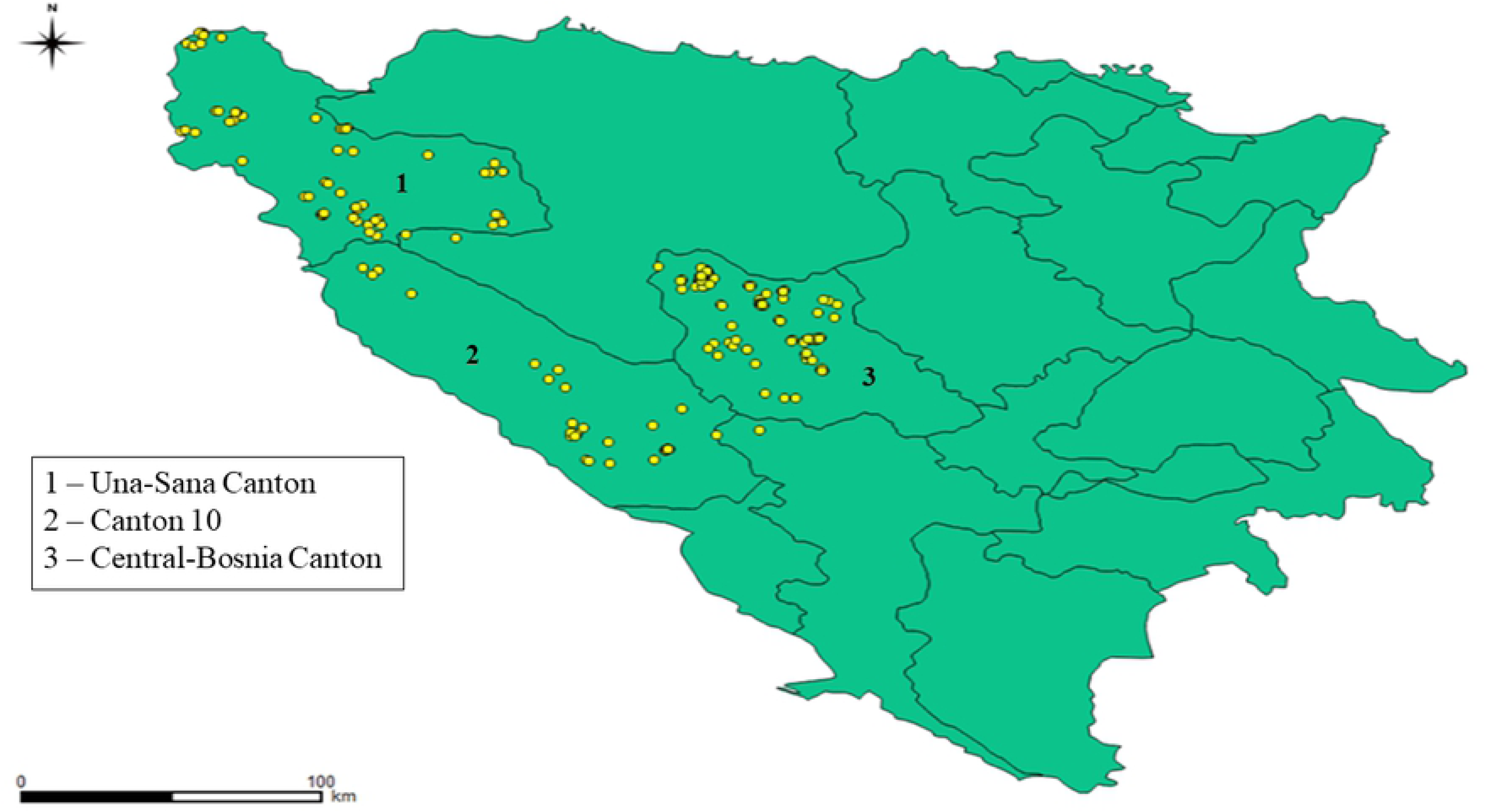
The distribution of 197 study farms located in three cantons of Bosnia and Herzegovina.

### Serological tests

Sera were analysed for antibodies against *C. abortus, C. burnetii, N. caninum*. The presence of antibodies against these pathogens was determined using the commercial Enzyme-Linked Immunosorbent Assays (IDEXX, Switzerland). Vaccination against *C. abortus, C. burnetii*, and *N. caninum* has not been implemented in BH, and hence, seropositive animals were considered to be naturally infected. Details of this serological survey are given in a former companion publication [6]. In the current study, we used the herd classification (presence or absence) for these agents based on prior publication.

### Reproductive outcomes (questionnaire)

The questionnaire was developed to collect information about management factors and reproductive underperformance in selected farms. Subsequently, the questionnaire was pre-tested in a small number of farms (n=5). After the pre-testing, the questionnaire was administered through a visit and an in-person interview with the farmer, made by the first author. The 40 questions were divided into three groups: socio-demographic information, management information, and information about reproductive performances in cattle. This questionnaire was also used in the previous paper [6], and may be found under this paper (https://bmcvetres.biomedcentral.com/articles/10.1186/s12917-018-1361-z). Supplementary rating questions were asked about the perceived primary constraints of the production. The pre-defined scale was based on Likert five-point scale (from “never” to “almost always”) [15].

### Variables

The primary reproduction outcomes defined in the present study were abortion, stillbirth, retention of placenta and metritis. Abortion was defined as any termination of the pregnancy after its confirmation by transrectal palpation or no return to oestrus. Transrectal palpation was done between 40 and 70 days after artificial insemination or natural breeding. In case of natural breeding, the date of such event was recorded by the farmer/owner. Stillbirth was recorded as a part of perinatal mortality. Perinatal mortality is defined as a death of the calf before, during or within 48h of calving at full term [16]. Retained placenta was considered to be any retention at 12 to 24h postpartum, while metritis was regarded as any condition with abnormal vaginal discharge, which was documented and treated by the veterinarian. We were not able to extract figures on the frequency of reproductive outcomes in each farm and had to rely upon information about farms’ and veterinarians written records in the period of five years.

The infectious outcomes defined in the current study were *C. abortus, C. burnetii* and *N. caninum* serostatus in the selected farms, based on results of our previous study [6]. Also, seropositivity to the listed reproductive infections was considered as an exposure factor for the occurrence of the aforementioned reproductive outcomes. Management-related factors were defined as potential predictors for both reproductive outcomes and reproductive diseases. A detailed description of all management-related factors investigated in this study is given in Tables 1-5. Further, the full list of management factors established as candidate variables is given in Table 6.

### Databases and statistical analyses

The questionnaire data were entered into a Microsoft Excel^®^ spreadsheet before inspected and merged with data from the previously reported serological survey (Softic et al., 2018). After cleaning data and initial analyses in Microsoft Excel^®^, using Pivot tables, data were transferred to Stata (Stata/SE 15 for Windows, StataCorp, College Station, TX, USA) for further statistical analyses.

Descriptive analyses and tabulation were done for each canton, and potential differences between cantons were assessed using a chi-square test for categorical variables and the Kruskal-Wallis test for continuous variables. After descriptive analyses, management-related factors were screened for their influence on abortion, stillbirths, retained placenta and metritis, as well as, infectious agents. As all reproduction and infection outcomes were coded as present/ absent (seropositive/ seronegative herd), a mixed-effects univariable logistic regression was used to screen management factors for potential statistical influence. Herd size effect on the occurrence of severe reproductive outcomes was expected, e.g. larger herds were more likely to report an abortion, stillbirth, metritis or retention of the placenta. Hence, an adjustment was made for herd size by including its natural logarithm (ln (herd size)) as an offset in the logistic model for the disease outcomes. This was not done for the infection outcomes, as this was based upon a more systematic sampling in the herds. Factors with a p-value less than or equal to 0.20 from univariable analyses were used as candidate variables for subsequent multivariable modelling.

The multivariable mixed-effects logistic models were established using the melogit procedure in Stata with canton as a random effect. Models were built using a backward selection procedure [15] with a set p-value less than or equal to 0.05 (5%) of the Likelihood Ratio test as exclusion criteria. After modelling all the management variables, the effect of herd infection status (*N. caninum, C. burnetii*, and *C. abortus*) were tested, by including these in the model and subsequently reducing the model using backward selection again. Notably, herd size variable was not adjusted and added as an offset in these models. Finally, all multivariable logistic models were assessed using the Hosmer-Lemeshow goodness-of-fit test and the area under the ROC curve (AUC) assessment on models based upon the *logistic* procedure [15].

The final multivariable models were then merged into a Structural Equation Model (SEM) using the *gsem* procedure in Stata. The structural equation modelling is a statistical framework used to model complex relationship between directly and indirectly observed variables [17]. In veterinary research, it is used as a primary statistical option [18] or as an additional technique based on previous statistical inference [19]. The SEM models, as available in the *sem* or *gsem* command in Stata, allows a confirmatory data analyses to be undertaken in the form of a path analysis, utilising all options built into Stata’s multilevel mixed-effects models. The initial model was constructed using the graphical *sembuilder* interface and then refined using the Stata command-language syntax. The model was built using the same strategy as the logistic model, but with a causal model specified. Additionally, the causal model was then specified graphically using the online software for Directed Acyclical Graphs (DAGitty) (Fig 2) [20]. As SEM allows an assessment of direct and indirect effects, we were able to reduce the standard logistic models further into more realistic causal models.

**Fig 2.**
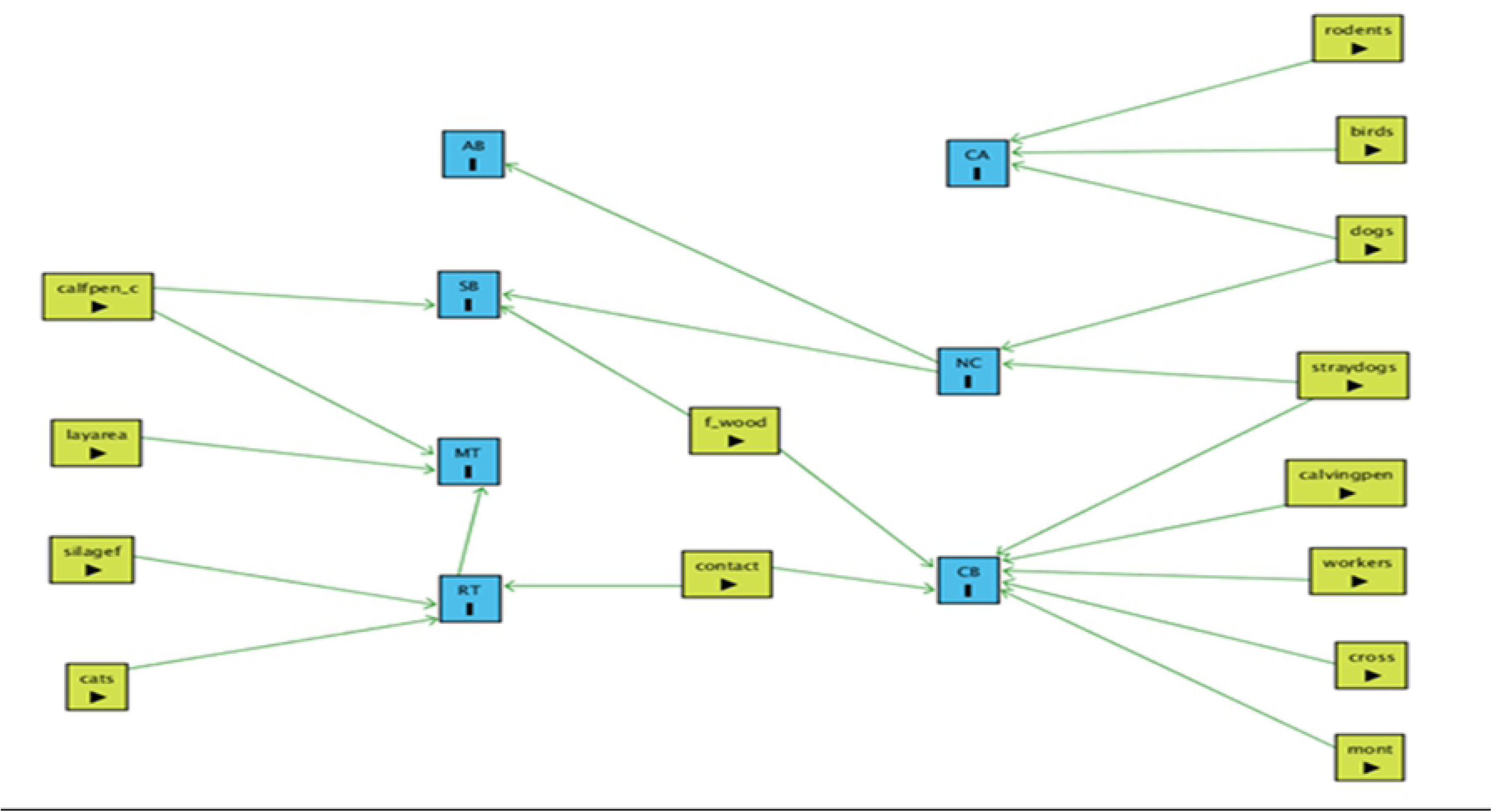
The suggested causal model based upon results from the confirmatory Structural Equation model. The graph was produced using the DAGitty software [20].

Assumed farm production constraints were analysed graphically using pivot functions in Microsoft Excel^®^ and visualised through a radar graph in Microsoft Excel^®^. Graphs were split according to herd size groups (quartiles), the variable found to determine the owners’ approaches to production constraints.

## Results

The study participants were distributed across three cantons of BH. The data were collected in a total of 19 municipalities and 94 local communities. There were a total of 66 (33.5%) farms in Una-Sana Canton, 35 (17.8%) farms in Canton 10 and 96 (48.7%) farms in Central Bosnia Canton. For the majority of farmers, cattle raising was not the only source of income but combined with other agricultural activities or non-agricultural employment. Altogether, 90% of selected farms had permanently employed workers, mainly family members, while a total of 10% farms used seasonally hired personnel.

Cattle in BH are usually housed during the winter period (December-April). During the rest of the year, cattle are allowed to graze on a daily basis and returned to the barn in the evening. Grasslands that cattle use for grazing surround the farm are displaced by a different distance from the farm. Farm facilities are usually located in the rural area, next to the family house.

### Housing and management

The distribution of farms by housing characteristics in three cantons in BH is shown in Table 1. In most of the visited farms (193/197, 98%), animals were tethered in the barns on wooden or concrete flooring.

**Table 1.** Distribution of 197 study farms by housing characteristics in three cantons in Bosnia and Herzegovina, given as median (range) for continuous or proportion (95% CI) for categorical variables.

| Variable | Region |  |  | Overall | p-value |
| --- | --- | --- | --- | --- | --- |
|  | Una-Sana Canton (n=66) | Canton 10 (n=35) | Central-Bosnia Canton (n=96) |  |  |
| Area of the barn | 100<br>(20-1600) | 100<br>(20-4000) | 40<br>(15-600) | 70 | <0.001 |
| Flooring type - wood | 28.8%<br>(19.1-41.0) | 57.1%<br>(40.2-72.5) | 84.4%<br>(75.6-90.4) | 60.9%<br>(53.9-67.5) | <0.001 |
| Flooring type - concrete | 92.4%<br>(82.9-96.9) | 77.1%<br>(60.1-88.3) | 40.6%<br>(31.2-50.8) | 64.5%<br>(57.5-70.9) | <0.001 |
| Presence of calving pen | 1.5%<br>(0.2-10.2) | 20%<br>(9.7-36.8) | 2.1%<br>(0.5-8.1) | 5.1%<br>(2.7-9.2) | <0.001 |
| Presence of calf pen | 93.9%<br>(84.8-97.7) | 94.3%<br>(79.4-98.6) | 75%<br>(65.3-82.7) | 84.8%<br>(79.0-89.2) | 0.001 |

A quarter of all farms used a combination of these two types of flooring. A total area of the barn’s facilities ranged from 20 to 4000 square meters, depending on herd size. Consequently, the laying area ranged from 1.5 to 5 m^2^ per animal. The majority (85%) of the farms had separate pens for calves, while a small number of the farms had separate calving pens. Substantial differences in farm management between cantons were observed. They are reflected in the farm size, number of animals per farm, and the breed composition. Also, there was a substantial difference in the presence of dogs and poultry on the farms in three cantons. On the other hand, there was no difference in feeding management between cantons.

In Table 2, basic information about herd sizes and management strategies are presented. The overall median number of herd size in the selected farms was six animals (range from 1 to 584). Despite the small number of cattle per farm, the majority (67.5%) of the farms used their stock as a source of replacement, while a limited number of the farms (5.1%) exclusively purchased animals for replacement. The breed composition of cattle in the selected farms was heterogeneous, with the Simmental as the most common breed observed in 76% of the farms. Distribution of farms by the presence of other domestic animals is shown in Table 3.

**Table 2.** Distribution of 197 study farms by a number of cattle and breed composition in three cantons in Bosnia and Herzegovina, given as median (range) for continuous or proportion (95% CI) for categorical variables.

| Variable | Region |  |  | Overall | p-value |
| --- | --- | --- | --- | --- | --- |
|  | Una-Sana Canton (n=66) | Canton 10 (n=35) | Central-Bosnia Canton (n=96) |  |  |
| No. of cattle |  |  |  |  |  |
| Herd size | 10 (2-101) | 11 (4-584) | 3.5 (1-150) | 6 | <0.001 |
| No. of cows per farm | 8 (2 - 62) | 10 (3-350) | 3 (0-112) | 5 | <0.001 |
| No. of heifers per farm | 2 (0-30) | 2 (0-234) | 0 (0-38) | 1 | <0.001 |
| No. Of calves per farm | 1 (0-25) | 1 (0-65) | 0 (0-94) | 1 | 0.003 |
| Breed composition |  |  |  |  |  |
| Cross breed | 51.5%<br>(39.4-63.4) | 54.3%<br>(37.6-70.1) | 69.8%<br>(59.8-78.2) | 60.9%<br>(53.9-67.5) | 0.043 |
| Holstein-Friesian breed | 21.2%<br>(12.9-32.9) | 17.1%<br>(7.8-33.7) | 8.3%<br>(4.2-15.9) | 14.2%<br>(10.0-19.9) | 0.06 |
| Simmental breed | 95.5%<br>(86.6-98.5) | 57.1%<br>(40.2-72.5) | 69.8%<br>(59.8-78.2) | 76.1%<br>(69.6-81.6) | <0.001 |
| Montafone breed | 4.5%<br>(1.5-13.4) | 62.9%<br>(45.7-77.3) | a | 12.7%<br>(8.7-18.2) | <0.001 |
| Red Angus breed | a | 2.9%<br>(0.4-18.3) | 1.0%<br>(0.1-7.2) | 1.0%<br>(0.3-4.0) | 0.395 |

a – no observation

**Table 3.**
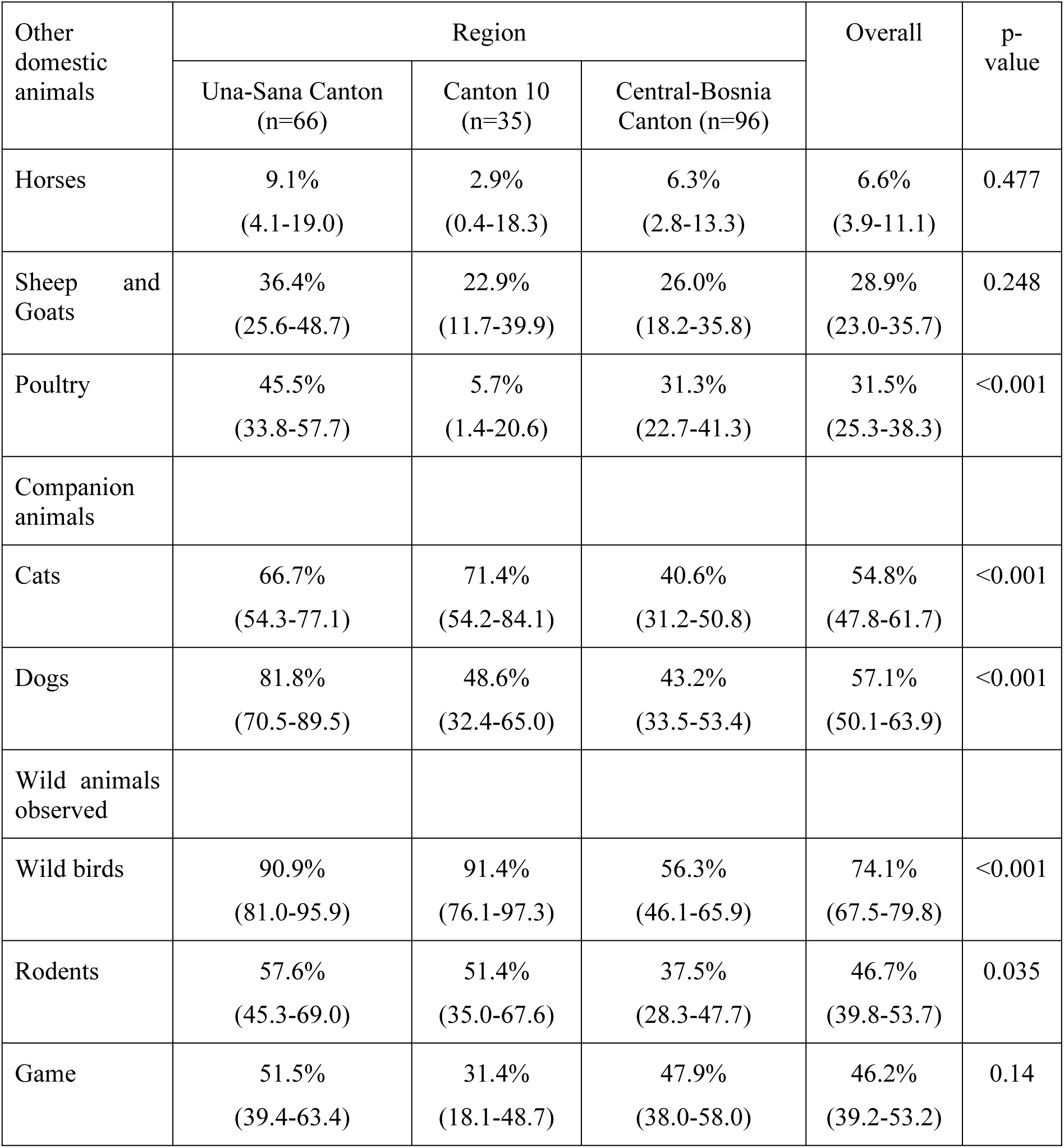
Distribution of 197 study farms by the presence of other species of animals in three cantons in Bosnia and Herzegovina, given as proportion (95% CI).

| Other domestic animals | Region |  |  | Overall | p-value |
| --- | --- | --- | --- | --- | --- |
|  | Una-Sana Canton (n=66) | Canton 10 (n=35) | Central-Bosnia Canton (n=96) |  |  |
| Horses | 9.1%<br>(4.1-19.0) | 2.9%<br>(0.4-18.3) | 6.3%<br>(2.8-13.3) | 6.6%<br>(3.9-11.1) | 0.477 |
| Sheep and Goats | 36.4%<br>(25.6-48.7) | 22.9%<br>(11.7-39.9) | 26.0%<br>(18.2-35.8) | 28.9%<br>(23.0-35.7) | 0.248 |
| Poultry | 45.5%<br>(33.8-57.7) | 5.7%<br>(1.4-20.6) | 31.3%<br>(22.7-41.3) | 31.5%<br>(25.3-38.3) | <0.001 |
| Companion animals |  |  |  |  |  |
| Cats | 66.7%<br>(54.3-77.1) | 71.4%<br>(54.2-84.1) | 40.6%<br>(31.2-50.8) | 54.8%<br>(47.8-61.7) | <0.001 |
| Dogs | 81.8%<br>(70.5-89.5) | 48.6%<br>(32.4-65.0) | 43.2%<br>(33.5-53.4) | 57.1%<br>(50.1-63.9) | <0.001 |
| Wild animals observed |  |  |  |  |  |
| Wild birds | 90.9%<br>(81.0-95.9) | 91.4%<br>(76.1-97.3) | 56.3%<br>(46.1-65.9) | 74.1%<br>(67.5-79.8) | <0.001 |
| Rodents | 57.6%<br>(45.3-69.0) | 51.4%<br>(35.0-67.6) | 37.5%<br>(28.3-47.7) | 46.7%<br>(39.8-53.7) | 0.035 |
| Game | 51.5%<br>(39.4-63.4) | 31.4%<br>(18.1-48.7) | 47.9%<br>(38.0-58.0) | 46.2%<br>(39.2-53.2) | 0.14 |

A total of 28.4% of the farms were exclusively cattle farms, while other farms practised combined keeping cattle with small ruminants (sheep and goats), poultry or horses. Also, more than half of the farms had companion animals, (dogs and cats). A total of 54.3% farmers have noticed stray dogs in and around their premises, especially in suburban areas. In 46.2% of farms, game (roe deer, wild boars, foxes) were observed within farms facilities or on grasslands. Stray dogs were included in the questionnaire as variable of interest as they are definitive host of *N. caninum* [8], and could directly and indirectly be included in the persistence of that agent in farms, while game can serve as a mechanical carriers of the studied infectious agents. A total of 48.7% of farmers reported herd-to-herd contact, mainly in the more densely populated villages close to cities. Rodents were observed in 46.7% of farms, and were more often found on larger farms that had a feed storage. Non-migratory wild birds were observed in a total of 74.1% of farms, with slightly difference between visited regions.

Depending on the season, roughage was based on grazing, feeding by cereals, industrial by-products and hay. Predominantly larger farms, i.e. farms with the greater number of animals (37.6%) have also used corn and grass silage. Water supply was ad-libitum usually coming from public or private sources (Table 4).

**Table 4.** Distribution of 197 study farms by characteristics for feeding and water supply in three cantons in Bosnia and Herzegovina, given as proportion (95% CI).

|  | Region |  |  | Overall | p-value |
| --- | --- | --- | --- | --- | --- |
|  | Una-Sana Canton<br>(n=66) | Canton 10<br>(n=35) | Central-Bosnia<br>Canton (n=96) |  |  |
| Feeding |  |  |  |  |  |
| Crops - cereals | 95.5%<br>(86.6-98.5) | 100% | 92.7%<br>(85.4-96.5) | 94.9%<br>(90.8-97.3) | 0.236 |
| Industrial by-products | 80.3%<br>(68.8-88.3) | 68.6%<br>(51.3-81.9) | 65.6%<br>(55.5-74.5) | 71.1%<br>(64.3-77.0) | 0.121 |
| Pasture | 81.8%<br>(70.5-89.5) | 88.6%<br>(72.8-95.7) | 91.7%<br>(84.1-95.8) | 87.8%<br>(82.4-91.7) | 0.168 |
| Hay | 100% | 100% | 97.9%<br>(91.9-99.5) | 99.0%<br>(96.0-99.7) | 0.345 |
| Silage | 74.2%<br>(62.2-83.5) | 31.4%<br>(18.1-48.7) | 14.6%<br>(8.8-23.2) | 37.6%<br>(31.0-44.6) | <0.001 |
| Water source |  |  |  |  |  |
| Public | 80.3%<br>(68.8-88.3) | 68.6%<br>(51.3-81.9) | 71.9%<br>(62.0-80.0) | 74.1%<br>(67.5-79.8) | 0.372 |
| Private | 10.6%<br>(5.1-20.8) | 8.6%<br>(2.7-23.9) | 13.5%<br>(8.0-22.1) | 11.7%<br>(7.9-17.0) |  |
| Both | 9.1%<br>(4.1-19) | 22.9%<br>(11.7-39.9) | 14.6%<br>(8.8-23.2) | 14.2%<br>(10.0-19.9) |  |

### Reproductive outcomes

The median age of animals at first artificial insemination was 15 months (range 11-24). Approximately 59% of farms used artificial insemination as a method of choice, while 41% used combined, natural breeding and artificial insemination. Almost all the visited farms have experienced some reproductive problems. Distribution of farms by the occurrence of reproductive problems is shown in Table 5. Reproductive problems such as abortion, stillbirth, metritis, and retained placenta were observed with substantial differences between cantons.

**Table 5.** Distribution of 197 study farms by the occurrence of reproductive problems in three cantons in Bosnia and Herzegovina, given as proportion (95% CI).

| Variable | Region |  |  | p-value |
| --- | --- | --- | --- | --- |
|  | Una-Sana Canton<br>(n=66) | Canton 10<br>(n=35) | Central-Bosnia<br>Canton (n=96) |  |
| Abortion | 69.7% (57.5-79.7) | 45.7%<br>(29.9-62.4) | 30.2% (21.8-40.2) | <0.001 |
| Stillbirth | 34.8% (24.2-47.2) | 37.1%<br>(22.7-54.3) | 14.6% (8.8-23.2) | 0.003 |
| Metritis | 45.5% (33.8-57.7) | 57.1%<br>(40.2-72.5) | 27.1% (19.1-39.9) | 0.003 |
| Retention of placenta | 72.7% (60.6-82.2) | 62.9%<br>(45.7-77.3) | 38.5% (29.3-48.7) | <0.001 |

Less observable disorders such as anoestrus, silent oestrus, and repeat breeder cows were recorded, but information on these was incomplete and are not included in the paper.

All potential factors listed in Tables 1-5 were tested in the univariable logistic regression. Table

6 displays the candidate variables showing a moderate association (p≤0.20) with severe reproductive outcomes and infectious agents that can lead to the reproductive failures.

**Table 6.** Candidate variables associated with the severe reproductive and infectious outcomes with P<0.2 from the univariable logistic regression, used as candidate variables for the multivariable logistic modelling.

|  | AB | MT | RT | SB | N. caninum* | C. burnetii* | C. abortus* |
| --- | --- | --- | --- | --- | --- | --- | --- |
| Region | x |  | x |  |  | x | x |
| Herd size (quartiles) | x |  |  | x | x | x |  |
| Crossbreed |  |  |  |  |  | x |  |
| Simmental breed |  |  |  | x |  |  | x |
| Montafone breed | x |  |  |  |  | x | x |
| Red Angus breed |  | x |  |  |  |  |  |
| Technical equipment |  |  |  |  | x | x | x |
| Workers |  |  | x |  | x | x |  |
| Area of the barn quartiles |  |  |  |  | x | x |  |
| Cleanliness degree | x |  |  |  |  |  |  |
| Wooden floor |  |  | x | x | x | x | x |
| Concrete floor |  |  | x | x | x | x | x |
| Laying area |  | x |  | x |  |  | x |
| Calving pen |  |  | x |  |  | x |  |
| Calf pen |  |  | x |  |  | x | x |
| Keeping calves in calf pen |  | x | x | x | x | x |  |
| Crops |  |  |  |  |  |  |  |
| By-product | x |  | x |  |  |  | x |
| Silage |  |  | x |  | x | x | x |
| Herd-to-herd contact |  | x | x | x | x | x | x |
| Horses |  |  |  |  | x |  |  |
| Small ruminants |  |  |  |  | x |  | x |
| Cats |  | x | x |  | x | x | x |
| Poultry | x |  |  |  |  |  |  |
| Birds |  | x |  | x | x | x | x |
| Hedgehog | x |  |  |  |  |  |  |
| Rodents | x |  |  |  |  | x | x |
| Straydogs | x | x |  | x | x | x | x |
| Dogs | x |  | x |  | x |  | x |
| R |  | x |  |  |  |  |  |
| N. caninum* | x |  | x | x |  |  |  |

AB - Abortions; MT- metritis; RT- retained placenta; SB- stillbirth; \* Seropositive herds

Table 7 shows the multivariable models for candidate variables that were associated with abortion, stillborn calves, retention of the placenta and metritis, as well as with the occurrence of *N. caninum, C. burnetii*, and *C. abortus* seropositive herds.

**Table 7.** Results from the final multivariable logistic models for the four reproduction outcomes and the infection outcomes

|  | Abortions | Stillbirth | Retention | Metritis | N. caninum* | C. burnetii* | C. abortus* |
| --- | --- | --- | --- | --- | --- | --- | --- |
| Rodents' presence | 2.1 (1.0-4.2);<br>p=0.047 |  |  |  |  |  | 4.0 (1.0-15.5);<br>p=0.044 |
| Dogs' presence | 2.2 (1.1-4.6);<br>p=0.033 |  |  |  | 3.00 (1.57-5.73); p=0.001 |  | 4.7 (1.5-14.6);<br>p=0.008 |
| N. caninum positive herd | 3.9 (1.9-7.9);<br>p<0.001 | 6.5(2.1-14.6);<br>p<0.001 |  |  |  |  |  |
| Hedgehogs' presence | 0.4 (0.2-0.8);<br>p=0.007 |  |  |  |  |  |  |
| Calving pen presence |  | 7.3 (1.4-38.4);<br>p=0.019 |  |  |  | 17.8 (2.4-134.0);<br>p=0.005 |  |
| Keeping calves in calf pens |  | Baseline |  | Baseline |  |  |  |
| In groups |  | 2.7 (1.1-6.5);<br>p=0.031 |  | 0.36 (0.16-0.83); p=0.016 |  |  |  |
| Workers |  |  |  |  |  | Baseline |  |
| Permanently employed |  |  |  |  |  | 0.2 (0.1-0.7);<br>p=0.011 |  |
| Silage - feeding |  |  | 3.1 (1.4-6.8);<br>p=0.004 |  |  |  |  |
| Herd-to-herd contact |  |  | 2.3 (1.1-4.6);<br>p=0.019 |  |  | 2.8 (1.0-7.8);<br>p=0.046 |  |
| Crossbreed |  |  |  |  |  | 0.4 (0.1-0.7);<br>p=0.049 |  |
| Montafon breed |  |  |  |  |  | 3.8 (1.2-11.8);<br>p=0.024 |  |
| Wooden floor presence |  | 2.3 (1.0-5.2);<br>p=0.048 |  |  |  | 0.2 (0.1-0.5);<br>p=0.001 |  |
| Stray dogs |  |  |  |  | 4.05 (2.12-7.72); p<0.001 | 3.8 (1.4-10.2);<br>p=0.01 |  |
| Cats' presence |  |  | 0.5 (0.3-1.0);<br>p=0.038 |  |  |  |  |
| Birds |  |  |  |  |  |  | 10.9 (3.6-33.5); p<0.001 |
| Laying area |  |  |  | 0.5 (0.3-0.8);<br>p=0.002 |  |  |  |
| Retention of placenta |  |  |  | 9.4 (4.0-22.2);<br>p<0.001 |  |  |  |

\* Seropositive herds

Table 8 and Fig 2 show how the models were transformed into a causally oriented SEM structure. As shown, the models were reduced in complexity due to the possibilities inherent in the SEM model to distinguish between direct and indirect causes. *N. caninum* was the only crucial infectious source for reproduction outcomes, while for some outcomes, environmental factors were important. Fig 2 illustrates our suggested causal structure, based on the results given in Table 8.

**Table 8.** Results from the confirmatory Structural Equation Model, based upon the multivariable logistic models found in Table 7.

| Outcome | Variable | OR | 95% CI | p-value |
| --- | --- | --- | --- | --- |
| Abortion (AB) | *N. caninum* seropositive herd (NC) | 4.3 | 2.2-8.4 | <0.001 |
| Stillbirth (SB) | *N. caninum* seropositive herd (NC) | 6.7 | 2.8-15.6 | <0.001 |
|  | Keeping calves in the calf pen (calfpen_c)-baseline |  |  |  |
|  | In groups | 2.6 | 1.1-6.4 | 0.032 |
|  | Wooden floor in the barn (f_wood) | 2.5 | 1.1-5.9 | 0.030 |
| Retention of the placenta (RT) |  |  |  |  |
|  | Feeding by silage (silagef) | 3.1 | 1.4-6.8 | 0.004 |
|  | Herd-to-herd contact (contact) | 2.3 | 1.2-4.6 | 0.019 |
|  | Cats' presence in the farm (cats) | 0.49 | 0.25-0.96 | 0.038 |
| Metritis (MT) | Retention of the placenta (RT) | 10.1 | 3.9-26.3 | <0.001 |
|  | Keeping calves in the calf pen (calfpen_c)-baseline |  |  |  |
|  | In groups | 0.36 | 0.15-0.82 | 0.016 |
|  | Laying area in the barn (layarea) | 0.46 | 0.28-0.78 | 0.004 |
| *N. caninum* seropositive herd (NC) | Dogs' presence in the farm (dogs) | 3.0 | 1.6-5.7 | 0.001 |
|  | Presence of the straydogs (straydogs) | 4.1 | 2.1-7.7 | <0.001 |
| *C. burnetii* seropositive herd (CB) | Presence of the calving pen in the barn (calvingpen) | 17.8 | 2.4-134.0 | 0.005 |
|  | Seasonally employed workers (workers)-baseline |  |  |  |
|  | Permanently employed workers | 0.18 | 0.05-0.68 | 0.011 |
|  | Herd-to-herd contact (contact) | 2.8 | 1.0-7.8 | 0.046 |
|  | Presence of the straydogs (straydogs) | 3.8 | 1.4-10.2 | 0.01 |
|  | Presence of the crossbred cows (cross) | 0.39 | 0.15-0.99 | 0.049 |
|  | Presence of the breed Montafone (mont) | 3.8 | 1.2-11.8 | 0.024 |
|  | Wooden floor in the barn (f_wood) | 0.17 | 0.06-0.49 | 0.001 |
| *C. abortus* seropositive herd (CA) | Dogs' presence in the farm (dogs) | 4.7 | 1.5-14.6 | 0.008 |
|  | Occurrence of the wild birds in the farm (birds) | 10.9 | 3.6-33.5 | <0.001 |
|  | Occurrence of the rodents in the farm (rodents) | 4.0 | 1.0-15.5 | 0.044 |

Outcomes: AB – Abortion; SB – Stillbirth; MT – Metritis; RT – Retained placenta; CA – C. abortus seropositive herd; NC – N. caninum seropositive herd; CB – C. burnetii seropositive herd.

Predictors: calfpen_c – Keeping calves in the calf pen individually (baseline) or in groups; layarea – Laying area in the barn; silagef – Feeding animals by silage; cats – Cats’ presence in the farm; f_wood – Wooden floor in the barn; contact – Herd-to-herd contact; rodents – The occurrence of the rodents in the farm; birds – The occurrence of wild birds in the farm; dogs – Dogs’ presence in the farm; straydogs – The occurrence of straydogs in and around farm’s facilities; calvingpen – The presence of the calving pen in the barn; workers – Seasonally (baseline) or permanently employed workers at the farm; cross – The presence of the cross breed at the farm; mont – The presence of the Montafone breed at the farm.

### Production constraints

Independent of herd size, the lack of market access due to production underperformance, and low purchasing prices of bovine products were the most common production constraints of all cattle farms in three cantons in BH (Fig 3).

**Fig 3.**
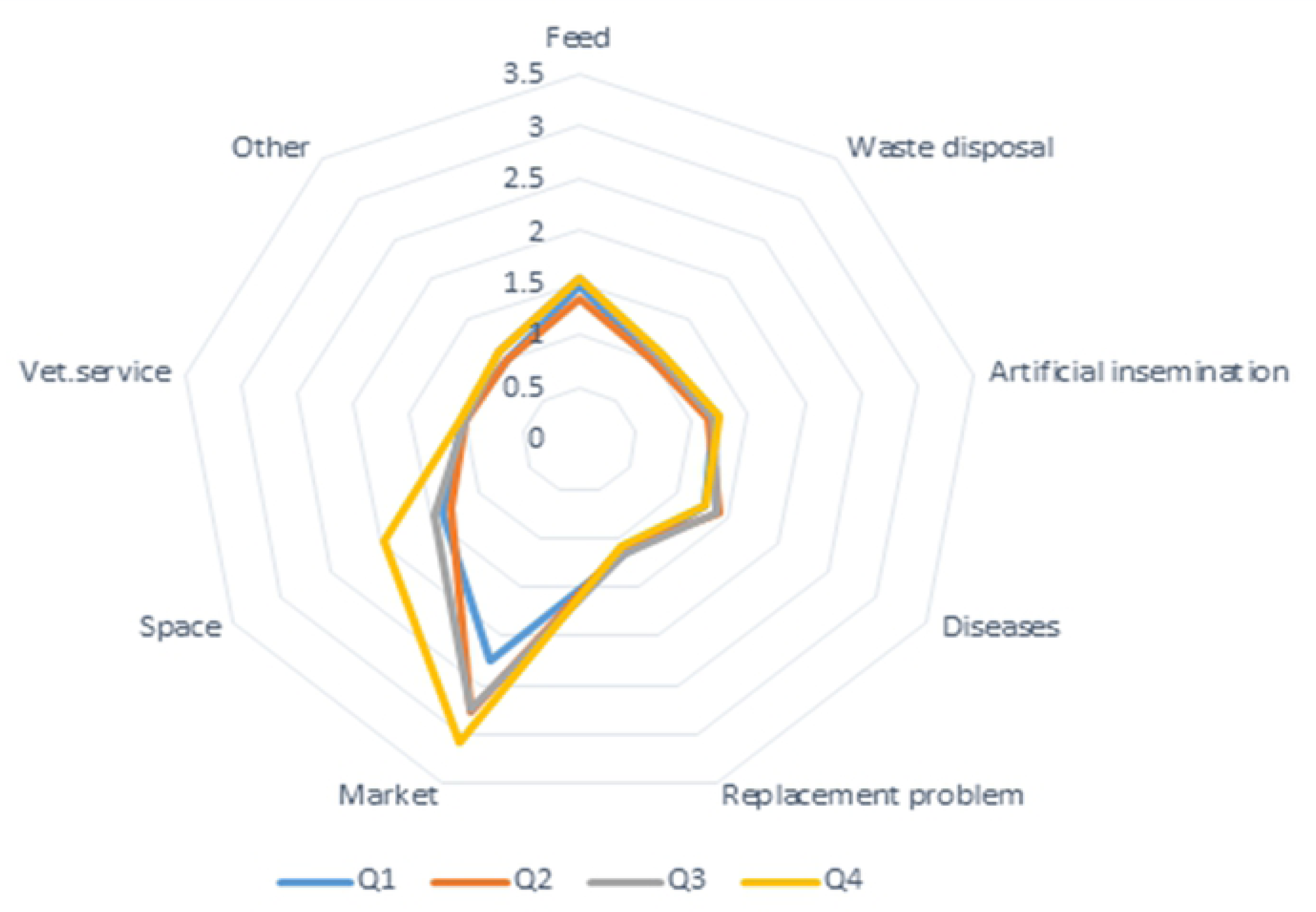
Main production constraints of 197 study farms in three cantons in BH shown on herd size quartiles.

## Discussion

Substantial differences in farm management between cantons were observed. These differences are mainly reflected in the herd size and structure. Farms located in the Una-Sana Canton and Canton 10 were mostly medium-sized to large, while farms located in the Central-Bosnia Canton were mostly small farms. Consequently, the number of animals per farm, and the breed composition were found significantly different. Also, the presence of poultry, companion animals (dogs and cats), wild birds, and rodents on the farms in three cantons were found significantly different. There were no differences between feeding practices between cantons.

In this study, the SEM was used as a confirmatory approach for assessing potentially unobserved direct associations between outcomes and predictors. Based on the results from the multivariable logistic regression, four different causal complexes were identified. In the following, we discuss the findings based on the SEM model in Table 8/ Fig 2.

### Abortion – Stillbirth – *N. caninum* herd seropositivity complex

One or more cases of abortion were observed in a total of 46.2% of the visited farms. Herds that were seropositive to *N. caninum* were more likely to report abortions compared to the herds that were seronegative to *N. caninum*. Several studies have observed an association between *N. caninum* seroprevalence in cattle herds and increased risk of abortion [21-23]. This is explained by an increased abortion risk because of vertical (latently infected dams) and horizontal transmission of *N. caninum*.

Results from the multivariable logistic regression showed that farms with dogs reported more abortions in cattle compared to farms with no dogs in their facilities. This may partially be explained by the *N. caninum* life cycle. As a definite host of the parasite, dogs participate in horizontal transmission through contamination of fodder by dog faeces [8]. Also, dogs might be an efficient vector of horizontal transmission due to permanent contact with cattle. Similar to our findings, other studies identified an association between farm dogs and bovine abortion as well as between the presence of dogs and the presence of *N. caninum* seropositive animals [21, 24]. Additionally, the occurrence of rodents or hedgehogs in and around farm facilities was associated with the abortions in cattle. The presence of these potential intermediate hosts of *N. caninum* might suggest that these animals could be important sources of infection for carnivores [8]. However, after SEM modelling these predictors disappeared as potential risk factors for abortion, leaving the seropositivity to *N. caninum* as the single explanatory variable.

A total of 25.3% of the visited farms reported stillbirth as a severe reproductive outcome. Herds that were seropositive to *N. caninum* were more likely (OR=6.5) to report stillbirth compared with the seronegative herds. Asmare et al. (2013) also found an association between abortion and/or stillbirth and individual animal level *N. caninum* seropositivity in Ethiopia [25]. Brickell et al. [26] reported an increased risk of perinatal mortality in *N. caninum* seropositive animals. The farms where the new-born calves were kept in groups reported more stillbirths than the farms where the new-born calves were kept individually. Additionally, farms with wooden flooring type had more (OR=2.5) stillborn calves compared to the farms with other flooring types. This could partially be explained by the calving management and calving assistance. There was an increased number of stillborn calves reported in the municipality of Livno (Canton 10) in 2015, which may partly be explained by inappropriate and unprofessional calving assistance, applied by farmers (Amel Murga, DVM, personal communication, April 18, 2015).

Antibodies against *N. caninum* were found in a total of 40.6% of the visited farms. Cattle in farms with the presence of dogs were more often (OR=3.0) seropositive compared to cattle in farms with no dogs. Similarly, farms that observed free-roaming dogs in and around their farm facilities had more seropositive (OR=4.1), compared to farms where no free-roaming dogs were observed. This finding is not surprising, as dogs are definitive hosts of *N. caninum.* Also, the farmer usually keeps dogs for many years, and it is often more than one dog per farm. In most epidemiological studies related to *N. caninum*, the presence of the farm dogs was found a risk factor for seropositivity in cattle [21-25]. However, caution is needed in the interpretation of these results. The Hosmer-Lemeshow goodness-of-fit test for the multivariable logistic regression model showed a lack of fit. It may be partially explained by the low number of groups in which the data were divided (n=4), while the recommendation is a minimum of 6 groups [15]. Another possible explanation could be that some cases of *N. caninum* herd seropositivity may result from exposure to risk factors and variables not included in this study. In summary, the SEM model allowed us to simplify and present a more appropriate causal model for this complex.

### Retention of placenta – Metritis complex

A total of 54.3% of the selected herds experienced retention of the placenta. Herds that fed predominately by silage had reported More retained placenta (OR=3.1) compared to herds that fed with a combined feed ration. This might be partially explained by nutritional demands in periparturient animals. Nutritional deficiencies, and consequently negative energy balance lead to metabolic disorders which represent a risk factor for the appearance of the retained placenta [27, 28]. Also, contact between herds seemed to be a substantial factor contributing to the appearance of retained placenta, possibly due to the transmission of infectious agents which can result in problems in the puerperium [7]. Additionally, such contact between animals in the field could plausibly mediate a part of the effects of uterine disease through social stress [29]. Herds that reported retained placenta had more (OR=9.4) metritis cases. The feature of the retained placenta as a predisposing factor for metritis is well documented [30, 31]. Manual removal of the retained placenta to prevent the source of infection is the dominant treatment among local veterinarians in the study area (Personal communication, January-August 2015). However, manual removal of an attached placenta may cause damage to the endometrium and could lead to bacterial invasion [32]. The retention-metritis complex, as shown by the SEM model, was associated with management factors, and we could not identify any infectious link here.

#### C. abortus

Farms that reported the presence of birds (pigeons, sparrows, swallows) had more (OR=11.0) animals seropositive to *C. abortus* compared to farms that did not report the presence of birds in their farm premises. Sachse et al. [33] found a few positive pigeons to *C. abortus* in their study of *C. psittaci*. Migration of these birds and their contact with dairy cattle or farm equipment could probably contribute to transmission of the agent, and consequently seropositivity in dairy cattle. Herds on the farms that noticed rodents (mice and rats) had more *C. abortus* (OR=4.0) compared to herds on the farms with no rodents in their farm facilities. Contrary to our findings, Sun et al. (2015) reported that management factors and presence of *Muridae* in farms were not associated with seropositivity of cattle to *C. abortus* [34]. Also, there was an association between dogs’ presence on the farm and seropositivity of dairy cattle. This complex was mainly linked to contact with other animals or environmental sources.

#### C. burnetii

Seropositivity to *C. burnetii* was associated with many farm management factors, contact between herds in the field, and presence of stray dogs in and around farm facilities. A possible explanation for this association is reflected in the indication that dogs (companion or feral) can be a potential reservoir species and source of infection for animals and humans [35]. Additionally, stray dogs as a population of homeless, abandoned dogs could be a particularly important source of infection for other animals and humans. Shapiro et al. (2016) observed that stray dogs from Aboriginal communities were more likely to be seropositive to *C. burnetii* than other population of dogs included in their study [36].

Herds that have contact with other herds on the common pastures had more seropositive to *C. burnetii* compared to isolated herds (OR=2.8). Contact with infected herds, contaminated equipment or biological material or other intermediate host seemed to be a possible explanation for this finding. Additionally, partial grazing system could favour seropositivity to *C. burnetii*. After re-housing in the winter period, seropositivity could be partially explained by exposure to aerosols or contaminated material [37].

Our suggested final causal model, depicted in Fig 2, joins a statistical model based upon SEM and a biological understanding of the reproductive diseases in question. The SEM model allowed us to conclude that there is a primary indirect effect of contact with dogs through *Neospora* infection, and we could also show the strict causality between placental retention and metritis, as the SEM models allow mediating effects in the model. A standard logistic model would not do this, as we would have to choose between correlated intervening variables. We did not assess model fits directly in SEM, as this was done in the original logistic models. The primary purpose of the SEM model was to reduce the complexity of the multivariable models and use SEM as a confirmatory technique to identify potential causal pathways.

The question of using appropriate statistical methods is a crucial part of epidemiological research. The development towards using mathematically more complex model has been fast, and today multivariable, multilevel regression has become a standard in analysing data from veterinary epidemiological studies, where hierarchical clustering often occurs. However, causal modelling has been used less. Most modern literature on causality and directed acyclical graphs, as used in this paper, is based upon the seminal book *Causality* by Judea Pearl [38]. In this book he argues well for using causal models including DAGs and either SEM (as used in this paper) or Bayesian Networks in causal inferences. Notably, these models are more efficient that traditional regression to deal with central topics as confounders and colliders, and also the complex problem of mediating variables. We support the idea that causal modelling is necessary in veterinary epidemiology, a science where we aim at suggesting interventions in populations and not only measure associations.

### Data quality

Recording of reproductive disorders is still at insufficient levels in BH. This was also reflected in our study, partially explained by the lack of an organised national cattle database. Farmers have limited skills in recognising a reproductive problem, and they seem to only on exceptional occasions seek professional assistance for contributing to data in the poor recording system. Reproductive outcomes such as abortion, stillbirth, retained placenta, and metritis are perceived as relatively severe, and the farmers can recognise the disease. Additionally, these outcomes affect (re)production, which also can motivate for consultancy. Subtler reproductive outcomes such as anoestrus and silent oestrus require more data and were not included in this study. Unfortunately, farms do typically not keep proper farm records, so we were not able to assess the frequency of these outcomes. Furthermore, farmers could overestimate or downscale the importance of specific factors investigated in this study. This was partially prevented by analysis of perceived production constraints.

### Production constraints

As the principal production constraint in this study, the respondents recognised inadequate and limited market access. This view is based on economic losses in their output. Such financial losses are categorised as direct, caused by relatively small and volatile purchasing prices of bovine products, and indirect, as a consequence of production underperformance. Furthermore, milk price volatility might be explained by criteria on milk quality (“Official Gazette BH” issue number 2205/2016). Changes in the composition of milk are leading to changes in its price. Also, milk price volatility could persist, following the permit for milk export to the EU and adapt to the external market laws. On the other hand, production underperformance could be partially explained by the occurrence of reproductive disorders, which result in direct loss of the offspring and indirect losses with reduced production in the recovering period.

### External validity of the findings

Although Western Balkans are at different stages of negotiations with the EU, apart from Croatia, they are faced with similar challenges. Mainly, their livestock farming is characterised by small and fragmented farms, with a small number of animals per farm. Such farm structure is inefficient and non-competitive either in export or in the domestic market. So far, a small number of farms have been commercialised and adapted to the requirements of the EU market. Also, traditional and outdated (extensive and semi-intensive) cattle rearing is still present in some areas of Western Balkans and represent an obstacle that is necessary to overcome. Given that most of these countries were members of the former Yugoslavia, the tradition of dairy farming and its management is more or less the same across them. One of the most important strategic points for improving dairy production is effective and timely dairy cattle reproduction. Previous studies have reported the presence of some reproductive agents in Croatia [39, 40], Serbia [41, 42] and Albania [43, 44]. Our findings bring a useful insight and can partially contribute to understanding the causality of management factors associated with reproductive underperformance and reproductive infections not only in BH but also in the Western Balkans.

## Conclusions

This study has demonstrated that management practices contributed to the occurrence of severe reproductive outcomes and reproductive infections. *N. caninum* seems to be an infectious agent that substantially contributed to the reproductive underperformance, particularly in inducing abortions and perinatal mortality. Also, there was a causal relationship between retention of placenta and metritis, which seems to be associated with poor managerial decisions. Our results illustrate the benefits of applying the SEM approach in elucidating potentially causal pathways in analysing complex epidemiological data.

## Acknowledgement

The authors express sincere gratitude to all study participants, veterinarians and contributors who helped in the development of this study.

## Author contributions

**Conceptualization:** AS, ES, KA

**Data curation:** AS

**Formal Analysis:** AS, ES

**Funding acquisition:** NF, ES

**Investigation:** AS

**Methodology:** AS, KA, EGG, ES, JG

**Project administration:** AS, NF, ES

**Supervision:** ES

**Validation:** EEG, ES, JG, NF

**Writing – original draft:** AS, ES

**Writing – review and analysis:** AS, KA, EEG, JG, NF, ES

